# An improved TEAD dominant-negative protein inhibitor to study Hippo YAP1/TAZ-dependent transcription

**DOI:** 10.1101/2024.10.03.615022

**Authors:** Briana Branch, Yao Yuan, Mariastella Cascone, Francesco Raimondi, Ramiro Iglesias-Bartolome

**Affiliations:** Laboratory of Cellular and Molecular Biology, Center for Cancer Research, National Cancer Institute, National Institutes of Health, Bethesda, Maryland, United States; Department of Cellular, Molecular, Developmental Biology and Biophysics Graduate Program, Johns Hopkins University, Baltimore, Maryland, United States; Laboratorio di Biologia Bio@SNS, Scuola Normale Superiore, Pisa, Italy

## Abstract

Hippo signaling is one of the top pathways altered in human cancer, and intensive focus has been devoted to developing therapies targeting Hippo-dependent transcription mediated by YAP1 and TAZ interaction with TEAD proteins. However, a significant challenge in evaluating the efficacy of these approaches is the lack of models that can precisely characterize the consequences of TEAD inhibition. To address this gap, our laboratory developed a strategy that utilizes a fluorescently traceable, dominant-negative protein named TEADi. TEADi specifically blocks the nuclear interactions of TEAD with YAP1 and TAZ, enabling precise dissection of Hippo TEAD-dependent and independent effects on cell fate. In this study, we aimed to enhance TEADi effectiveness by altering post-transcriptional modification sites within its TEAD-binding domains (TBDs). We demonstrate that a D93E mutation in the YAP1 TBD significantly increases TEADi inhibitory capacity. Additionally, we find that TBDs derived from VGLL4 and YAP1 are insufficient to block TAZ-induced TEAD activity, revealing crucial differences in YAP1 and TAZ displacement mechanisms by dominant-negative TBDs. Structural differences in YAP1 and TAZ TBDs were also identified, which may contribute to the distinct binding of these proteins to TEAD. Our findings expand our understanding of TEAD regulation and highlight the potential of an optimized TEADi as a more potent, specific, and versatile tool for studying TEAD-transcriptional activity.

## MANUSCRIPT

Hippo signaling is downstream from mechanosensing and receptor-mediated pathways that regulate cell fate, helping cells balance cell growth and differentiation ^1-3^. At its core, Hippo signaling is a kinase cascade that results in the cytoplasmic retention and inactivation of YAP1 and its paralog TAZ (also known as *WWTR1*). YAP1 and TAZ interact with numerous effectors, although their regulatory and oncogenic functions are mainly mediated by binding the family of TEAD transcription factors (TEAD1 to 4) ^4-6^.

The recognized oncogenic properties of YAP1 and TAZ have solidified them as primary targets for developing inhibitors to treat cancer and other hyperproliferative diseases, mainly focused on blocking YAP1 and TAZ interaction with TEAD. However, the lack of models to precisely characterize the consequences of TEAD inhibition is a significant challenge in studying the efficacy of this approach. In addition, YAP1 and TAZ interact with numerous other proteins in the cytoplasm and the nucleus, making it difficult to discern TEAD-dependent and independent events.

Several tools are available to characterize the disruption of YAP1/TAZ-TEAD binding and its use for cancer treatment, including dominant-negative proteins, peptides that block YAP1-TEAD interactions, and drugs that inhibit TEAD activity. One of the first strategies used was to block TEAD function by overexpressing truncated TEAD proteins lacking the DNA-binding domain, acting as dominant-negative forms ^5,7^. A similar approach has been to utilize fusion proteins of the TEAD4 transcription factor with the repression domain of Drosophila Engrailed (Tead4EnR) or the transcriptional activation domain of herpes simplex virus VP16 (Tead4VP16) ^8^. Small peptide inhibitors, such as the Super-TDU peptide ^9^, have been derived from the VGLL4 and YAP1 TEAD binding domains (TBDs) to block YAP1-TEAD interaction. Verteporfin, the first drug characterized to inhibit TEAD, works by binding YAP1 and blocking its interaction with TEAD ^7^. IAG933 is another drug that directly binds TEAD and prevents its interaction with YAP and TAZ ^10^. Recent studies have established that TEAD proteins require palmitoylation for activation ^11,12^, leading to the development of small molecule inhibitors that target the TEAD lipid site ^13-15^. While each of these tools offers unique advantages, they lack one or more essential elements, including traceability, efficient inhibition of both YAP1 and TAZ or specific blockage of TEAD nuclear events without affecting YAP1 and TAZ interaction with cytoplasmic effectors.

To address these limitations, our laboratory developed a complementary strategy that utilizes a fluorescently traceable, dominant-negative protein named TEADi that specifically blocks the nuclear interaction of TEAD with YAP1 and TAZ ^16^ (Fig. 1A). TEADi offers several advantages for studying YAP1/TAZ-TEAD dependent transcription, including easy visualization, rapid and simple inhibition of TEAD, and specific blockage of nuclear events mediated by both YAP1 and TAZ without affecting their structural or cytoplasmic functions ^16^. TEADi has been used to elucidate Hippo signaling across diverse contexts ^17-28^. Despite TEADi advantages, a key challenge of dominant-negative proteins is the need for high expression levels and strong binding to effectively outcompete and displace endogenous interacting partners of the target protein. Hence, improving TEADi stability and affinity for TEAD could improve its inhibitory capacity, allowing for effective TEAD inhibition at lower expression levels.

**Figure 1:**
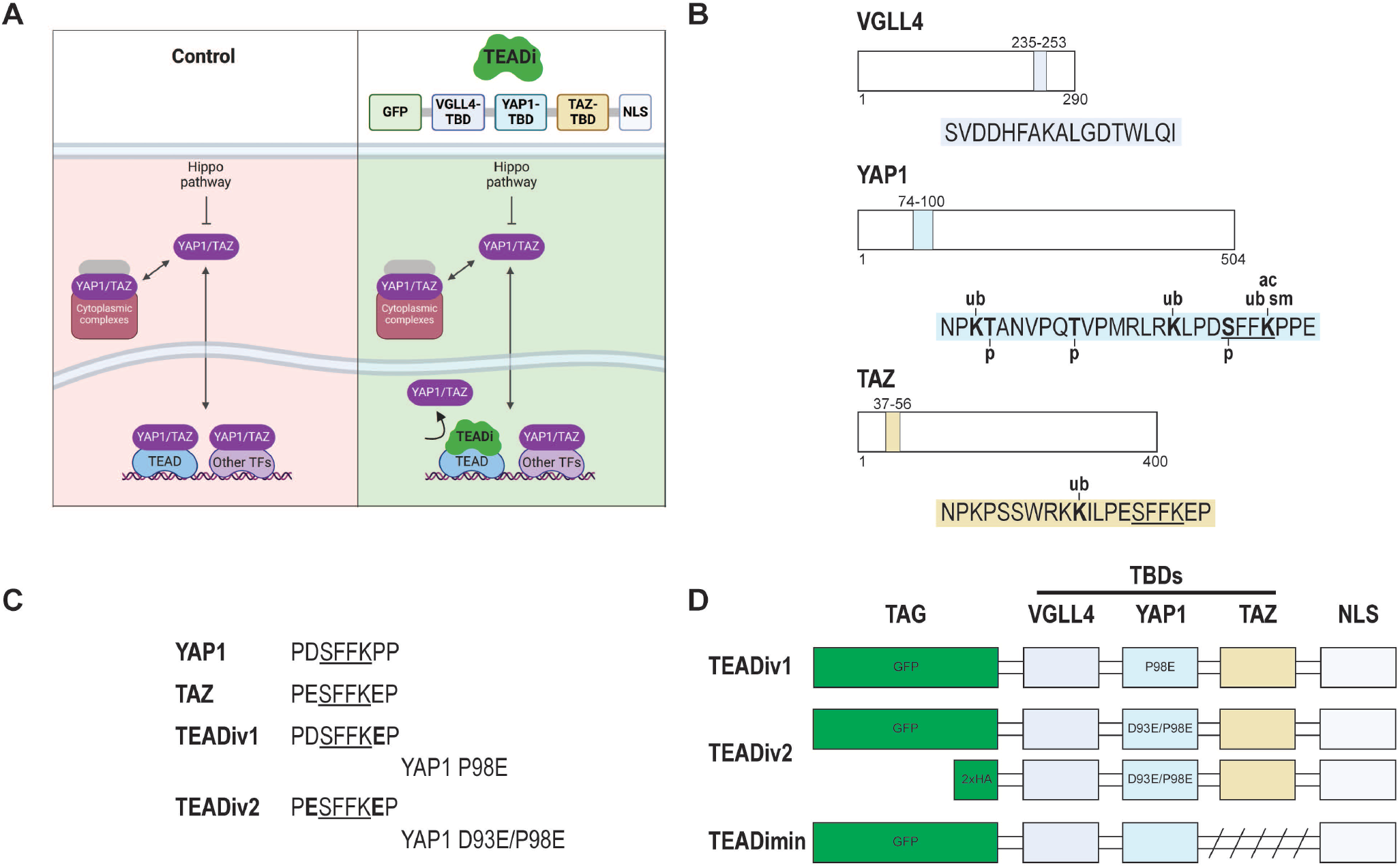
Analysis of post-transcriptional modifications in TEAD binding domains and development of a next-generation TEAD inhibitor peptide. **A**. Model showing Hippo signaling and how TEADi disrupts YAP1/TAZ binding to TEAD transcription factors, specifically in the nucleus. The building blocks of TEADi are shown. TBD= TEAD-binding domain, NLS= nuclear localization signal. **B**. Detail of the TBDs that compose the TEAD inhibitor. Colors indicate the stretch of the amino acid sequence from each protein found in TEADi. Reported post-transcriptional modifications in TBDs are indicated (www.phosphosite.org^29^). p= phosphorylation, ub= ubiquitination, sm= SUMOylation, ac=acetylation. The homologous site in YAP1 and TAZ that binds TEAD is underlined. **C**. Detail of the homologous site in YAP1 and TAZ that binds TEAD and mutations introduced in TEADiv1 and TEADiv2. **D**. Depiction of the building blocks for constructions used in this study (not at scale).

TEADi is built upon the TEAD binding domains (TBDs) of VGLL4, YAP1, and TAZ ^16^ (Fig. 1A). Since YAP1 and TAZ nuclear localization and interaction with TEAD are tightly regulated by phosphorylation and ubiquitination ^2^, we hypothesized that altering post-transcriptional modification sites in the TEADi TBDs could enhance its stability and inhibitory capacity. Analysis of post-transcriptional modifications in TEADi TBDs indicated the presence of phosphorylation, ubiquitination, and SUMOylation sites, particularly in the YAP1 TBD (www.phosphosite.org ^29^) (Fig. 1B). Phosphorylation S94 in YAP1 is known to disrupt its binding to TEAD transcription factors ^30,31^ and it is in a region essential for YAP1-TEAD interaction, indicating that preventing S94 phosphorylation could improve binding between these proteins. However, it has been shown that mutating the S94 residue leads to a loss of YAP1-induced TEAD activity and TEAD binding ^5,30^. A different way to prevent S94 phosphorylation would be to mutate surrounding residues to avoid kinase binding. Since the homologous site in TAZ (S51) is not phosphorylated (Fig. 1B, underlined) and this region has a high affinity for TEAD ^32^, we changed aspartic acid 93 in the YAP1 TBD to glutamic acid (D93E), mimicking the TAZ sequence (Fig. 1C). The original TEADi had already a P98E substitution in the YAP1 TBD that increases its affinity to TEAD ^16,32^ (Fig. 1C). We named the new version TEADiv2 and the original as TEADiv1 (Fig. 1C and D). We also constructed a version of TEADi without the TAZ TBD that resembles the Super-TDU inhibitor peptide ^9^ (minimal TEADi or TEADimin) and a TEADi version with an HA tag instead of GFP to characterize potential effects from the fluorescent tag (Fig. 1D).

Transduction of cells with plasmids corresponding to each construct indicates that they are well expressed at the predicted molecular weight (Fig. 2A). All TEADi constructs reduce the expression of the YAP1/TAZ-TEAD target CYR61, while, in general, they do not affect the expression of TEAD, YAP1 or TAZ (Fig. 2A). In agreement with the effect of TEADiv1 ^16^, coimmunoprecipitation experiments confirmed that TEADiv2 blocks the interaction of YAP1 and TAZ with TEAD transcription factors (Fig. 2B). To compare the efficacy of TEADi versions we measured TEAD-induced transcriptional activity by co-transfecting cells with different doses of DNA for each dominant-negative protein in the presence of YAP1 or TAZ and a luciferase reporter composed of eight repeats of the TEAD binding site (8x-TEAD-Luc ^33^). TEADiv2 showed increased inhibition of YAP1-induced TEAD activity at lower doses than TEADiv2 (Fig. 2C). In contrast, both proteins showed a similar profile of inhibition towards TAZ (Fig. 2D). These results are in agreement with the modifications of TEADiv2 being in the YAP1 TBD (Fig. 1C). We also observed that TEADi without a TAZ TBD (TEADimin) showed a decrease inhibition towards TAZ (Fig. 2E), indicating that the presence of the TAZ TBD enhances the activity of TEADi towards this protein and pointing towards potential differences in YAP1 and TAZ displacement from TEAD by the dominant negative TBDs.

**Figure 2:**
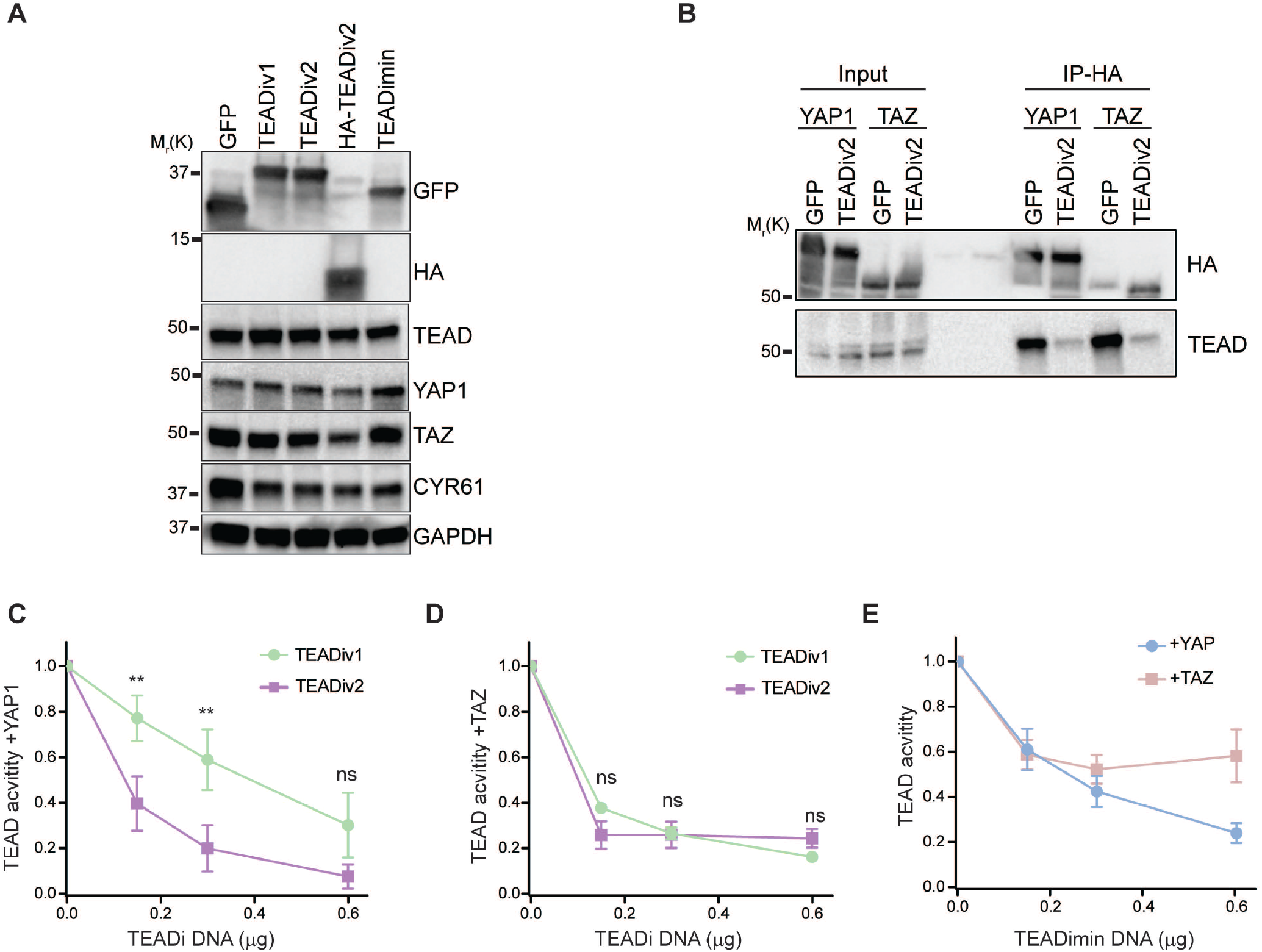
TEADiv2 shows enhanced TEAD inhibition towards YAP1-TEAD activity. **A**. Western blot showing the expression of TEAD inhibitors in HEK293 cells 24hs after transfection. Unless otherwise indicated, TEADiv1 and TEADiv2 refer to the GFP-tagged version. **B**. Western blot of coimmunoprecipitation experiments showing that TEADiv2 reduces TEAD interaction with YAP1 and TAZ. **C-D**. Dose curve comparing TEADiv1 and TEADiv2 TEAD inhibition with 8x-TEAD-Luc reporter in HEK293 cells overexpressing YAP1 (C) or TAZ (D). n=3 independent experiments with technical duplicates. ns=p>0.05, **=p<0.005, paired t test. Mean ± SEM is shown. **E**. Dose curve comparing TEADimin TEAD inhibition with an 8x-TEAD-Luc reporter in HEK293 cells overexpressing YAP1 or TAZ. n=3 independent experiments with technical duplicates. Mean ± SEM is shown.

To measure the effectivity of TEADiv2 against each member of the TEAD family, we utilized TEAD transcription factors fused to a Gal4 DNA-binding domain ^5^. As shown in Fig. 3A, TEADiv2 efficiently blocked YAP1 and TAZ-induced activity for all TEAD family members. Finally, we measured TEAD inhibition by a GFP or HA-tagged TEADiv2 protein using the 8x-TEAD-Luc reporter. Our results show that YAP1 and TAZ-induced TEAD activity was similarly blocked by both tagged versions of TEADiv2 (Fig. 3C), indicating that the efficiency of the dominant-negative protein is independent of the tag.

**Figure 3:**
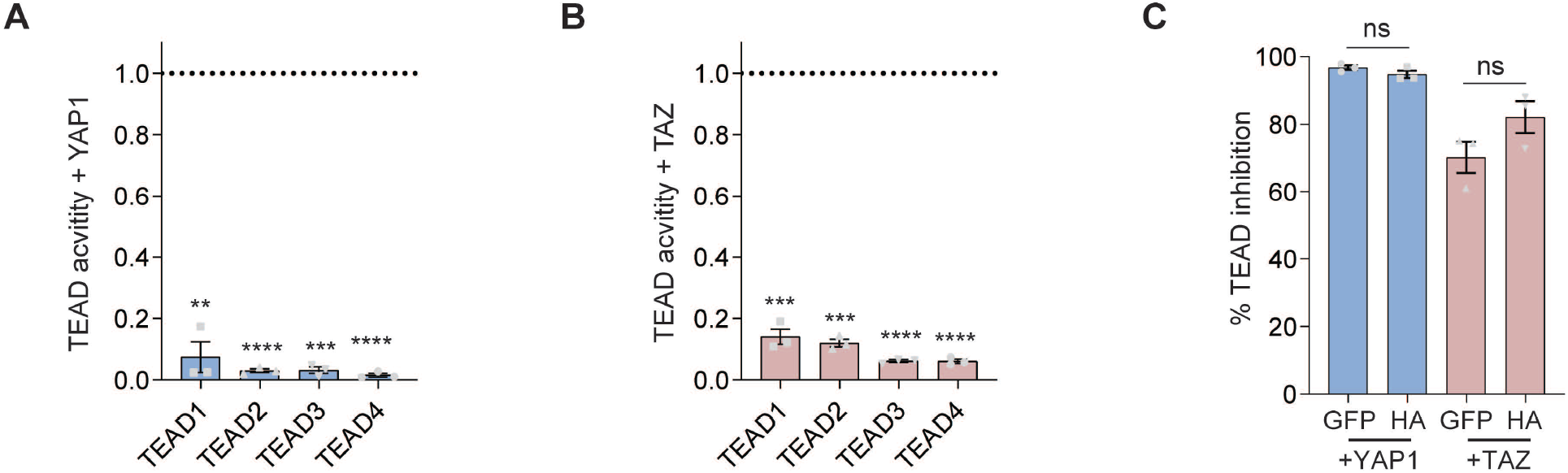
TEADiv2 inhibits YAP1 and TAZ TEAD activity for all TEAD family members, and its activity is independent of the tag. **A-B**. Graphs showing TEAD activity inhibition by TEADiv2. Transcriptional activity of TEAD was measured by luciferase assay in HEK293 cells transduced with TEADiv2 or GFP, each corresponding TEAD fused to the Gal4 DNA binding domain, UAS-luciferase, and either YAP1 (A) or TAZ (B). TEAD activity is reported as the fold decrease in TEADi compared with GFP control (dotted line indicates activity in GFP condition). n=3 independent experiments with technical duplicates. **=p<0.005, ***=p<0.001, ****=p<0.0001, t test comparing GFP vs TEADi. Mean ± SEM is shown. **C**. Comparison of GFP-tagged and HA-tagged TEADiv2 with 8x-TEAD-Luc reporter in HEK293 cells overexpressing YAP1 or TAZ. TEAD activity is reported as percentage inhibition over the activity of cells transfected with GFP instead of TEADi. n=3 independent experiments with technical duplicates. ns=p>0.05, t test. Mean ± SEM is shown.

Next, we performed structural predictions with AlphaFold-Multimer ^34,35^ to understand the potential differences between TEADiv1 and TEADiv2. We observed that, in isolation, TEADiv2 adopts an alternative conformation with respect to TEADiv1 (Fig. 4A). We then used RosettaRelax ^36^ to refine and score predicted structures, with lower energy values translating to a more favorable conformation ^37^. TEADiv2 is predicted to be more energetically stable, since its structural models (considering the top10 predictions) are associated with more negative energies compared to TEADiv1 (Fig. 4B). This also correlates in part with a local structuring of TEADiv2 N- and C-terminals, predicted to have higher α-helix content (Figure 4A). Prediction of the complex between TEADi and TEAD4 suggested different 3D conformations for the TEADiv2 complex (Fig. 4C), correlated with overall lower energies of the complexes (Fig. 4D). The predicted structural differences between TEADis are interesting considering that the only difference between TEADiv1 and TEADiv2 is the change of aspartic acid 93 in the YAP1 TBD to glutamic acid (D93E). Our modeling indicates that this D93E substitution does not affect the local structure in the mutation site but translates into a different predicted peptide conformation. Despite the similarities between aspartates and glutamates, it has been proposed that for intrinsically disordered regions aspartates support extended structures and helical caps, while glutamates support increased helicity, which is necessary for folding and binding ^38^. These differences in the properties of aspartates and glutamates to drive structure suggest that the increased activity of TEADiv2 could be partly due to a more stable conformation of the inhibitor resulting from the D93E change in the YAP1 TBD.

**Figure 4:**
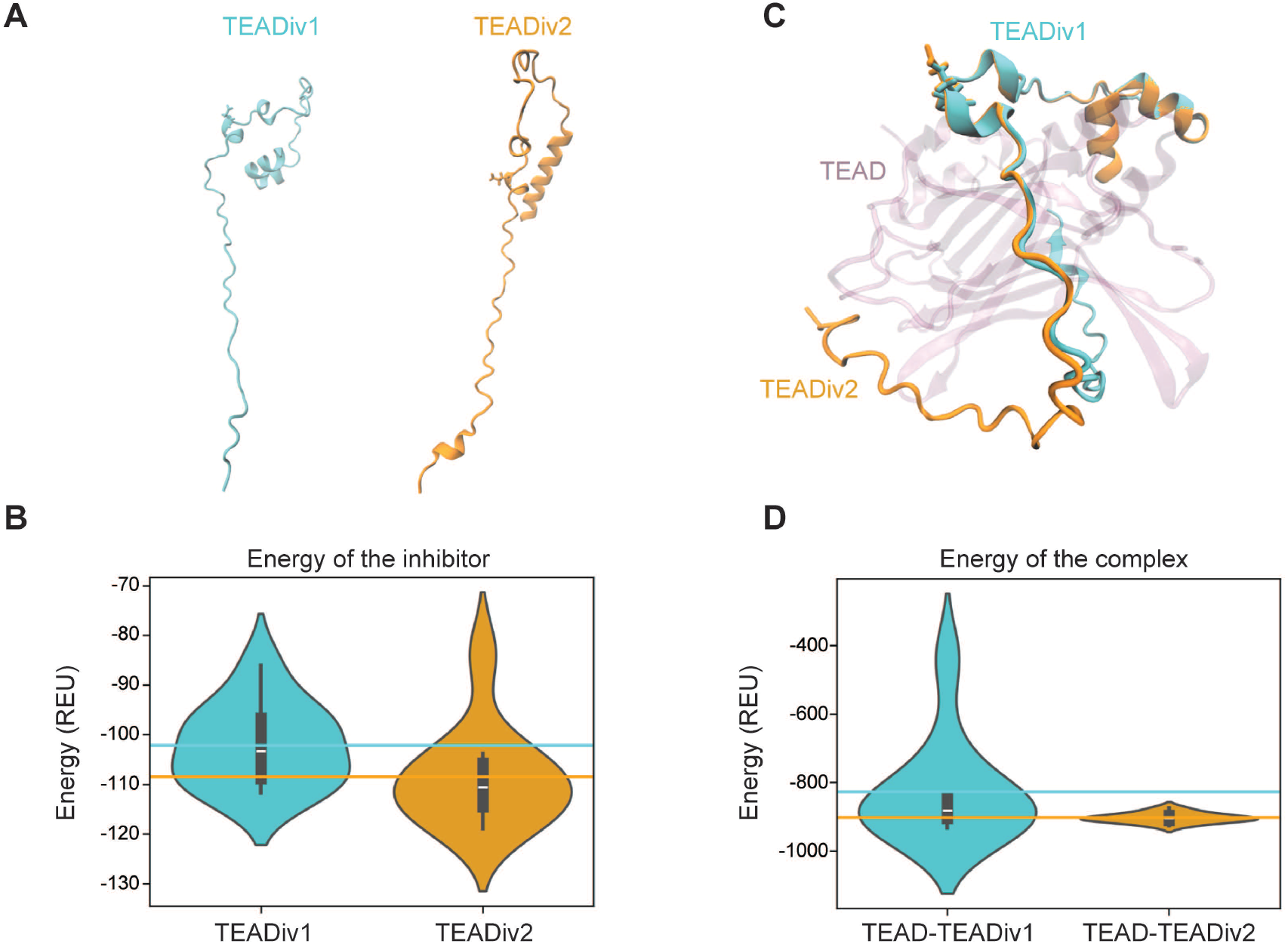
TEADiv2 is structurally more stable and has increased affinity for TEAD. **A**. Predicted 3D conformations of isolated TEADiv1 and TEADiv2 via AlphaFold-Multimer v2.3.2. **B**. Violin plot depicting the energy distribution -in Rosetta energy units (REU)-of the top 10 predicted models of TEADiv1 and TEADiv2. Violin plots show density of data and lines indicate the corresponding average. **C**. Predicted 3D modeling of TEADiv1 and TEADiv2 interaction with TEAD4 (purple). **D**. Violin plot depicting the Rosetta energy units (REU) of the top 10 predicted models of TEADiv1 and TEADiv2 in complex with TEAD. Violin plots show density of data and lines indicate the corresponding average.

Overall, our study reveals new insights into the regulation of TEAD activity by YAP1 and TAZ and highlights the potential for improving dominant-negative TEAD inhibitors. We demonstrate that altering post-transcriptional modification sites can enhance the activity of peptides based on TBDs, making them more effective in blocking TEAD-mediated transcriptional activity. We also find that the VGLL4-YAP1 TBDs are insufficient to inhibit TAZ-induced TEAD activity, indicating differences in how YAP1 and TAZ are displaced from TEAD by dominant-negative TBDs. Our results further demonstrate that TEADi effectively inhibits the activity of all four TEAD transcription factors driven by YAP1 and TAZ. Finally, we also identify structural differences in the TBDs of YAP1 and TAZ that may underlie their differential binding and affinity for TEAD. Our work highlights the potential of TEADiv2 as a more effective, specific, and versatile tool for studying TEAD transcriptional activity.

## MATERIALS & METHODS

### DNA constructs and cell culture

TEADiv1 construct has been described before ^16^. TEADiv2 and TEADimin were constructed using gBlocks Gene Fragments (Integrated DNA Technologies) downstream of GFP or 2 sequences of the hemagglutinin A tag (2xHA) into a pCEFL vector. Human YAP1 plasmid was constructed in gBlocks Gene Fragments (Integrated DNA Technologies) using sequence optimized YAP1 (NM_001282101) with an N-terminal 3xHA tag followed by a 3xGS linker and cloned into pCEFL. pCEFL vector has an EF-1α promoter and was a gift from Silvio Gutkind. 8x-TEAD-Luc (pGL3b 8xGTIIC-luciferase) was a gift from Stefano Piccolo ^33^ (Addgene plasmid #34615). pcDNA3-HA-TAZ ^39^ and pCMX-GAL4-TEAD1 to 4 ^5^ constructs were a gift from Kunliang Guan (Addgene plasmids 32839, 33108, 33107, 33106, and 33105 respectively). pGL4.31[luc2P/Gal4UAS/Hygro] was purchased from Promega (catalog #C9351). HEK293 cells were cultured at 37 °C in the presence of 5% CO_2_. These cells were obtained from AddexBio and cultured in DMEM (Sigma-Aldrich Inc) supplemented with 10% fetal bovine serum (FBS) (Sigma-Aldrich Inc) and antibiotic/antimycotic solution (Sigma-Aldrich Inc).

### Luciferase assays

Luciferase assays were performed in HEK293 cells. To measure pan-TEAD activity, cells were plated in poly-d-lysine coated 24-well plates and transfected 24 h after plating. Cells were co-transfected overnight with 8x-TEAD-Luc (0.3 µg cm^−2^) plus the indicated DNA constructs. Unless stated otherwise, the DNA amount used was: GFP (0.3 µg cm^−2^), TEADis (0.3 µg cm^−2^), YAP1 (0.1 µg cm^−2^), TAZ (0.1 µg cm^−2^). The following day, cells were harvested, and luciferase activity was measured using a Dual-Glo Luciferase Assay Kit (Promega) and a Microtiter plate luminometer (SpectrMax iD3, Molecular Devices LLC). To measure individual TEAD family member activity, cells were co-transfected with pCMX-GAL4-TEAD1 to 4 constructs (0.3 µg cm^−2^), pGL4.31[luc2P/Gal4UAS/Hygro] Vector (0.2 µg cm^−2^) plus the indicated DNA constructs at 0.3 µg cm^−2^ and processed as described above. Luciferase normalization was performed in every case by co-transfecting a Renilla luciferase vector (0.01 µg cm^−2^).

### Immunoprecipitation

For immunoprecipitation (IP), the anti-HA tag antibody was used (Covance; clone no. 16B12; catalog no. MMS-101R; 1:100). Proteins were extracted in IP lysis buffer (50 mM Tris-Cl pH 7.5, 150 mM NaCl, 10% glycerol, 2 mM MgCl_2_, and 1% NP40) supplemented with a complete protease inhibitor cocktail (Roche, #6538304001). Protein extracts were subjected to centrifugation at max speed for 10 min, and then supernatant protein was quantified. Equal amounts of protein were incubated in a rotator at 4 °C overnight with primary antibody. Then, protein extracts were incubated in a rotator for two hours at 4 °C with Dynabeads Protein G (Invitrogen, #10004D). Finally, beads were washed 4 times with IP lysis buffer, and proteins were eluted with denaturing loading buffer for Western blot analysis.

### Immunoblot analysis

Cells were lysed by sonication at 4 °C in lysis buffer (50 mM Tris-HCl, 150 mM NaCl, 1 mM EDTA, 1% Nonidet P-40, 0.2% sodium deoxycholate, 0.1% SDS, 1 mM DTT) supplemented with complete protease inhibitor cocktail (Roche, #6538304001) and phosphatase inhibitors (PhosSTOP, Sigma-Aldrich, #4906837001). Equal amounts of total cell lysate proteins were subjected to SDS-polyacrylamide gel electrophoresis and transferred to PVDF membranes. Protein visualization was performed using the following antibodies: anti-GAPDH (Cell Signaling; clone no. 14C10; catalogue no. 2118; 1:2000), anti-GFP (Cell Signaling; clone no. D5.1; catalog no. 2956; 1:2000), anti-HA tag antibody (Cell Signaling; clone no. C29F4; catalog no. 3724; 1:1000), YAP1 (Cell Signaling; clone no. D8H1X; catalog no. 14074; 1:1000), TAZ (Cell Signaling; clone no. V386; catalog no. 4883; 1:1000), CYR61 (Cell Signaling; clone no. D4H5D; catalog no. 14479; 1:1000), and Pan-TEAD (Cell Signaling; clone no. D3F7L; catalog no. 13295; 1:1000). Secondary HRP-conjugated antibodies used were: Pierce peroxidase goat antimouse IGG (H + L) (ThermoFisher, catalog no. 31432; 1:2500), and Multi-rAb HRP-Goat Anti Rabbit Recombinant Secondary Antibody (H + L) (Proteintech, catalog no. RGAR001; 1:30000). Secondary antibody was incubated at room temperature for 2 h. Bands were detected using a ChemiDoc Imaging System (Bio-Rad) with Clarity Western ECL Blotting Substrates (Bio-Rad) according to the manufacturer’s instructions.

### Structural predictions

we employed AlphaFold-Multimer v2.3.2 ^34,35^ with jackhmmer to produce 25 structures of each complex (TEAD-TEADiv1 and TEAD-TEADiv2) and 25 of each inhibitor in its isolated form (TEADiv1 and TEADiv2). We inputted the following sequence stretches for TEADiv1 and TEADiv2, encompassing the VGLL4, YAP1 and TAZ domains:TEADiv1:SVDDHFAKALGDTWLQIGDPPVATNPKTANVPQTVPMRLRKLPDSFFKEPEGDPPVATNPKPSSWRKKILPESFFKEP;TEADiv2:SVDDHFAKALGDTWLQIGDPPVATNPKTANVP QTVPMRLRKLPESFFKEPEGDPPVATNPKPSSWRKKILPESFFKEP. For TEAD4, we inputted the human canonical sequence from Uniprot (Uniprot Accession: Q15561). For each modeling run, the top-10 scoring structures were then relaxed using Rosetta Relax version 2021.16.61629 (fast relax protocol) ^36^, employing backbone restraints (flag: “constrain_relax_to_start_coords”). From the outputs of the relax, we then extracted the total energies (in Rosetta Energy Units - REU) and structures. We generated violin plots using the “violinplot” function of the seaborn (v0.13.2) python library. 3D representation was generated using VMD (v1.9.4) ^40^.

### Statistics

All analyses were performed in triplicate or greater and the means obtained were used for t-tests. Statistical analyses, variation estimation and validation of test assumptions were carried out using the Prism 9 statistical analysis program (GraphPad).

## ACKNOWLEDGMENTS

This research was supported by the Intramural Research Program of the National Institutes of Health, National Cancer Institute, Center for Cancer Research (ZIA BC 011763).

